# HIPSD&R-seq enables scalable genomic copy number and transcriptome profiling

**DOI:** 10.1101/2023.10.09.561487

**Authors:** Olga Lazareva, Jan-Philipp Mallm, Milena Simovic-Lorenz, George Philippos, Pooja Sant, Urja Parekh, Linda Hammann, Albert Li, Umut Yildiz, Mikael Marttinen, Judith Zaugg, Kyung Min Noh, Oliver Stegle, Aurélie Ernst

## Abstract

Single-cell DNA-sequencing (scDNA-seq) enables decoding somatic cancer variation. Existing methods are hampered by low throughput or cannot be combined with transcriptome sequencing in the same cell. We propose HIPSD&R-seq (**HI**gh-through**P**ut **S**ingle-cell **D**na and **R**na-seq), a scalable yet simple assay to profile low-coverage DNA and RNA in thousands of cells in parallel. Our approach builds on an accessible modification of the 10X Genomics platform for scATAC and multiome profiling. In applications to human cell models and primary tissue, we demonstrate the feasibility to detect rare clones and we combine the assay with combinatorial indexing to profile over 16,000 cells.

## Background

Advances in single-cell DNA sequencing (scDNA-seq) technologies have enabled exploring new dimensions of genomic diversity that are not accessible by bulk sequencing. scDNA-seq methods have been instrumental in the fields of somatic mutation and mosaicism [1–6], organ development [7–11], germ cell mutation and fertility [12–16], epigenetic regulation and genome organisation [17–23] and cancer research [1,2,24–27]. Established protocols conduct targeted sequencing, i.e. focus on individual mutations of interest [28–34], or consider low-coverage genome-wide sequencing (scDNA-seq) for the identification of copy number variants or somatic alterations (CNVs and CNAs, respectively) [35–39]. Here, we focus on the latter, which has essential implications in the context of precision oncology and resistance to cancer treatment.

The first scDNA-seq protocols have been extended and improved in different ways, to increase coverage and reduce bias (the latter through amplification-free workflows for example) [26,40–42] and to allow for parallel profiling of DNA and RNA from the same cells [36,37,42–44]. Most recently, with the advent of combinatorial indexing, the feasibility of scaling scDNA-seq to tens of thousands of cells in a single experiment has been demonstrated [46]. However, what is lacking to date are schemes that render high-throughput scDNA-seq accessible and reproducible across laboratories. Commercially available technologies such as the 10x platforms have had tremendous impact on RNA and epigenome profiling, yet there is no equivalent for the scalable profiling of genomic variation in single cells. In fact, the lack of accessible scDNA-seq protocols has stimulated the development of computational solutions to infer coarse grained CNAs from RNA abundance profiles, a strategy that is recognized to have lower resolution and accuracy [47,48], yet achievable using commercial single-cell RNA sequencing (scRNA-seq) platforms [47,49–51].

Key challenges of current protocols for scDNA-seq [2,26,40–44,44,52–56] include low throughput and/or difficulty to establish these methods elsewhere, for instance due to the need for specialised equipment [57]. As compared to scRNA-seq, which has been democratised by commercially available systems [58–67], scDNA-seq and parallel single-cell DNA and RNA sequencing (scDNA/RNA-seq) are lagging behind. For parallel analyses of copy-number variation and transcriptomes from the same cells, pioneering studies have focused on relatively small numbers of cells to demonstrate methods like DR-Seq (13 to 33 cells) [40], Simul-seq (8 to 10 cells) [68], scTrio-seq (25 cells) [37], SIDR-seq (74 cells) [44] or scONE-seq (86 cells) [69] to at most hundreds of cells for G&T-seq (220 cells) or DNTR-seq (230 cells) [36,70]. sci-L3 recently provided a proof-of principle in cell lines for the analysis of DNA and RNA of thousands of cells [71], with scalable combinatorial indexing as a major advance. A notable exception is the approach proposed developed in parallel by Peter Sims and colleagues [72], which shares some of the characteristics of our method. However, our approach and in particular the DNA analyses to resolve clonal heterogeneity provide a number of advantages, and we also achieve a four-fold higher yield, providing an ultra-high throughput. Here, we propose **HI**gh-through**P**ut **S**ingle-cell **D**na and **R**na-seq (HIPSD&R-seq), a scalable yet simple assay, which offers high-throughput scDNA-seq from several hundreds to thousands of cells, coupled with the benefits of an established workflow that builds on the commercial 10x Genomics platform. We compare HIPSD&R-seq to alternative scDNA-seq protocols and to CNV inference based on scRNA-seq, finding substantial benefits in resolution and accuracy. Finally, we demonstrate the utility of the throughput provided by HIPSD&R-seq in the context of a spike-in experiment to detect rare clones, and we apply the assay to human fibroblasts and to medulloblastoma patient-derived xenografts.

## Results

We have developed HIPSD&R-seq, a modular experimental workflow that allows for implementing scalable single-cell DNA and optionally RNA sequencing from the same cells (**Fig. 1**). Core to our approach is the repurposing of existing scATAC-seq and multiome protocols to perform high-throughput scDNA-seq (**Fig. 1**). Briefly, Tn5 has been a versatile tool for many sequencing applications, including ATAC-seq [18], RNA library preparation [73–75], 3D chromatin structure [76–79] and also DNA-seq [36,40,44]. We utilised highly concentrated Tn5 *in situ* by fixing cells mildly with formaldehyde, depleting nucleosomes with SDS followed by the transposition reaction in still intact nuclei. Combined with the 10X Genomics scATAC-seq kit to encapsulate cells, this modification provides the means to achieve medium-throughput single-cell DNA-seq (HIPSD-seq, > 5,000 cells per sample). The highly concentrated Tn5 replaced the supplied enzyme by 10X to ensure high integration rates across the genome. The same modification of the genome tagmentation can also be combined with the 10X multiome kit to enable parallel scDNA-seq and RNA sequencing (HIPSD&R-seq). Finally, HIPSD-seq can also be combined with an additional combinatorial indexing step [46] to achieve ultra-high throughput scDNA-seq (sci-HIPSD-seq, > 10,000 cells) (**Methods**). Data from all three protocols can be processed using the standard Cell Ranger pipelines. We have further developed computational strategies to aggregate read data across cells, which are beneficial when estimating CNV profiles from sparse high-throughput scDNA-seq protocols such as HIPSD&R-seq (**Methods**). Briefly, to mitigate coverage sparsity, we have developed a metacelling workflow, which adapts existing strategies from scRNA-seq to scDNA-seq. By aggregating reads data from genetically similar cells, this approach greatly improves read coverage (to approximately at least 0.02x) and uniformity while preserving rare clones.

**Figure 1.**
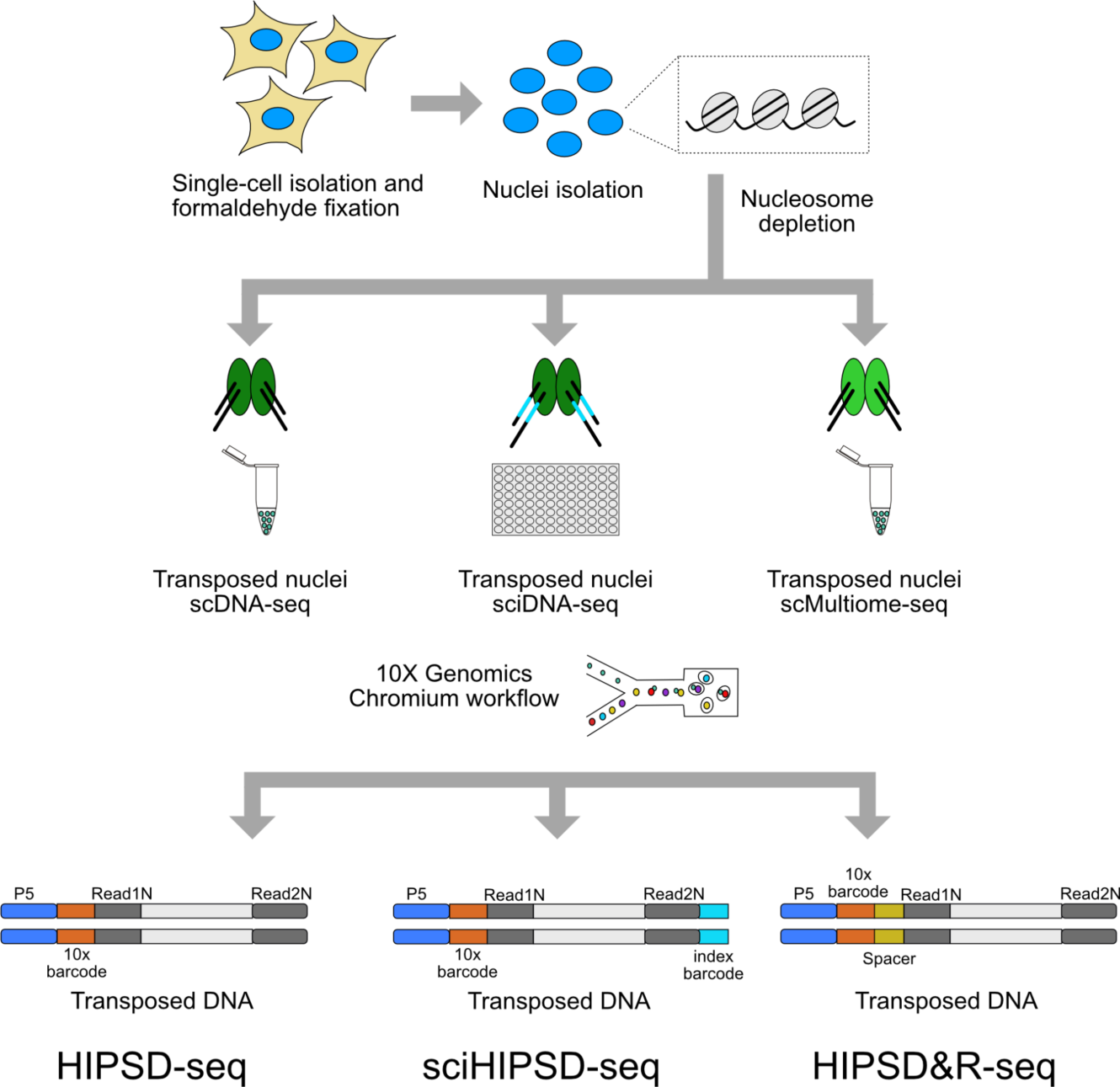
Schematic representation of the experimental approach for three connected protocols based on a modular modified 10x Genomics workflow - HIPSD-seq, sci-HIPSD-seq and HIPSD&R-seq. Formaldehyde fixed single-cell suspensions are used for nuclei isolation, followed by SDS-based nucleosome depletion. The nucleosome-depleted genomic regions are randomly tagmented using the transposase Tn5 (HIPSD-seq and HIPSD&R-seq) or barcoded Tn5 (sci-HIPSD-seq). Transposed nuclei are processed using the 10x Genomics Chromium single cell ATAC workflow (HIPSD-seq and sci-HIPSD-seq) or the multiome workflow (HIPSD&R-seq) to generate single-cell DNA and gene expression (RNA) libraries for sequencing. Combining scDNA-seq with an additional combinatorial indexing step thanks to the barcoded Tn5 leads to an ultra high throughput (sci-HIPSD-seq).

First, to assess the quality of HIPSD-seq libraries, we processed nuclei from human fibroblasts with high genome instability [80]. HIPSD-seq yielded more than 5,000 high-quality cells per sample (**Suppl. Table 1**, **Suppl. Fig. 1**). We assessed canonical quality control metrics, including read uniformity and the total DNA sequence coverage (**Suppl. Fig. 1**). These data showed that while HIPSD-seq on individual cells yields lower coverage than existing scDNA-seq methods, the combination of HIPSD-seq with metacelling yields data that are comparable to existing sparse scDNA-seq methods in terms of coverage (**Suppl. Fig. 1**). As a reference for desired quality characteristics, we benchmarked HIPSD-seq with data from the discontinued Chromium Single Cell CNV assay [81], obtained from the same PDX sample (**Methods**, for more details see also Parra et al [82]). Additionally, we compared quality of transcriptome obtained from HIPSD&R-seq to Chromium Single Cell Gene Expression assay on the same cells (for more details see also Parra et al [82] and **Suppl. Fig. 1**).

We next applied HIPSD&R-seq to conduct low-coverage DNA and RNA sequencing of cells from a patient-derived xenograft of a medulloblastoma with high genome instability. The RNA component of HIPSD&R-seq recovered a median number of over 1,800 transcripts per cell (**Suppl. Table 1**), which is in line with data from conventional multiome assays [36,44]. Motivated by the uptake of computational approaches to infer CNAs from scRNA-seq data, we compared CNV estimates from the DNA component of HIPSD&R-seq to results obtained using inference from the corresponding RNA profiles (using Numbat, Methods) [47]. Qualitatively, the CNV states from HIPSD&R-seq offered markedly higher resolution (**Fig. 2A**). We also assessed the concordance of the CNV estimates with the corresponding bulk whole-genome CNV profiles from the same sample. This comparison revealed substantially higher consistency for HIPSD-seq compared to CNV inference from the matched RNA (**Fig. 2B**), in particular in the regime of smaller size CNVs. Overall, HIPSD&R-seq resolves clonal heterogeneity that results from CNVs and provides matched transcriptome profiles.

**Figure 2.**
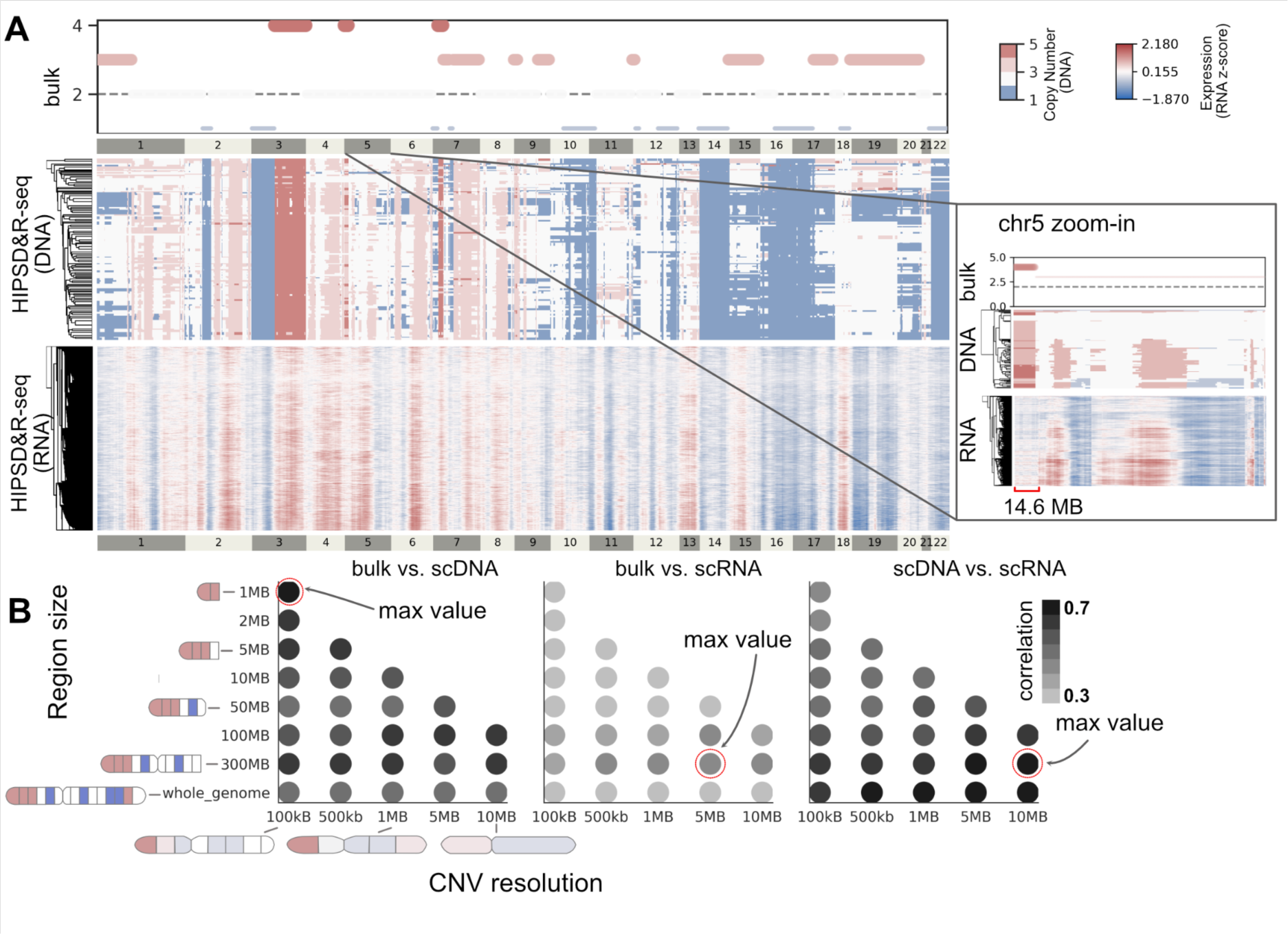
HIPSD&R-seq shows close correspondence to the reference bulk data (patient-derived xenograft from a medulloblastoma). **(A)** From top to bottom: CNV estimates derived from bulk WGS data (top), CNV estimates derived from HIPSD&R-seq in 101 meta cells (middle), CNV estimates derived from the HIPSD&R-seq RNA component (bottom, numbat window-smoothed expression). DNA-based CNV estimated obtained using HMMcopy; RNA-based CNV estimated derived using numbat. Right: Zoom-in view on chromosome 5, depicting a representative region where CNV estimates derived from RNA failed to capture a localised amplification event (marked in red). **(B)** Consistency analysis of CNV estimates derived from bulk DNA and from the DNA and RNA components of HIPSD&R-seq. Shown are average correlation coefficients for pairs of alternative methods, varying the CNV quantification resolution (x-axis) and the size of genomic regions for accession correlation (y-axis). From left to right: bulk DNA versus HIPSD&R-seq DNA, bulk DNA versus HIPSD&R-seq RNA, HIPSD&R-seq DNA versus HIPSD&R-seq RNA.

Finally, we set out to test ultra-high throughput scDNA-seq using sci-HIPSD-seq, which is particularly suitable to identify rare sub populations of cells. We performed a spike-in experiment by mixing 1% of nuclei from fibroblasts from a male donor (LFS087) to a nuclei suspension from a female individual (LFS041). The CNV profiles from the combined sample revealed a sub population that is consistent in frequency with the 1% prevalence of the spike-in sample (**Fig. 3**). Assessment of the concordance with the bulk profiles from either patient samples (LFS087 and LFS041) confirmed the identity of both sub populations, thus demonstrating the ability to detect rare cell populations using sci-HIPSD-seq. As our HIPSD-seq methods are easy, high-throughput and affordable, they pave the way to applications in cancer genomics, especially in the context of chromosome instability and treatment resistance.

**Figure 3.**
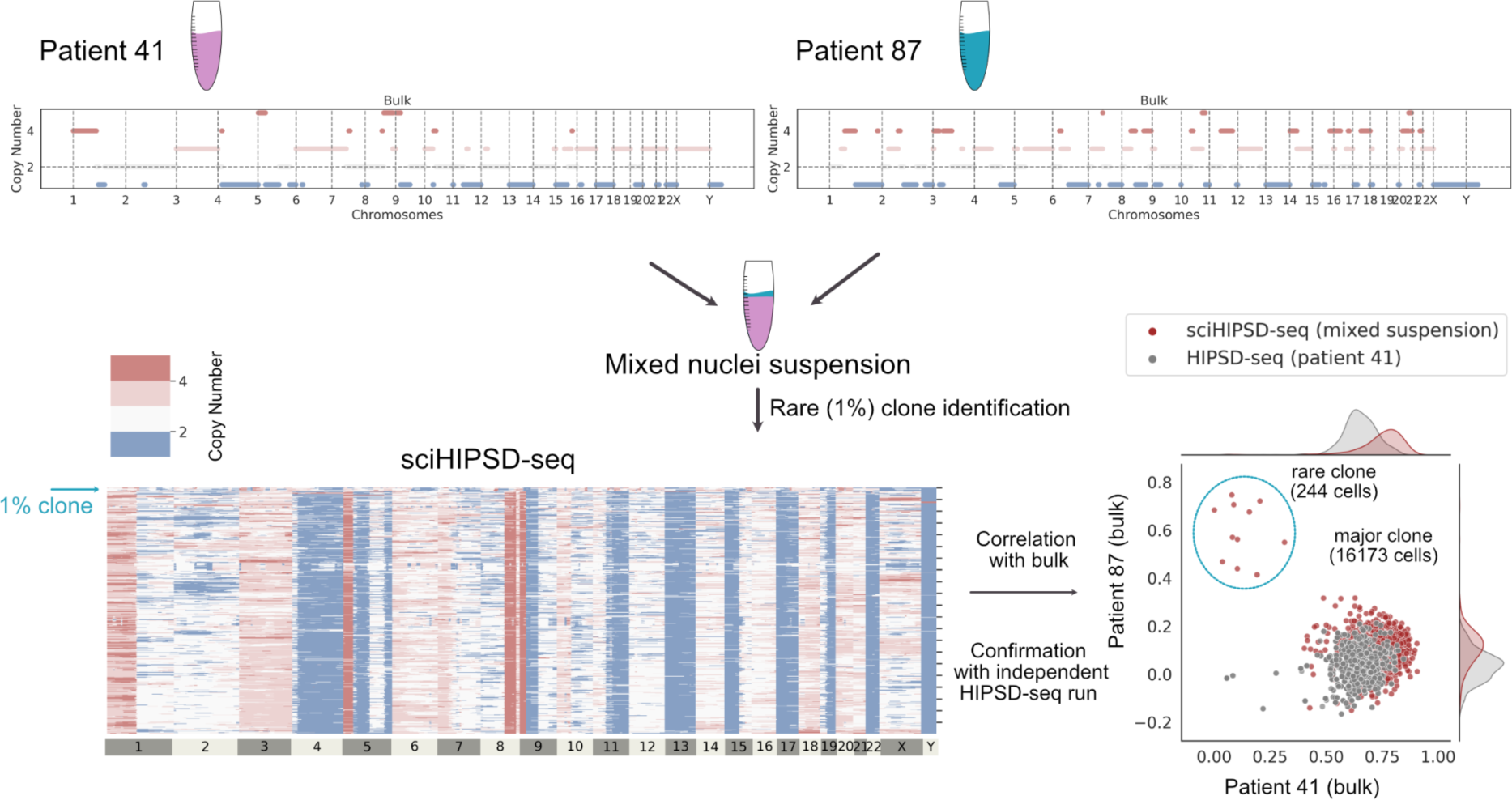
sciHIPSD-seq identifies a rare clone. Top: Bulk profiles for patient 87 (1% spike-in) and patient 41 (99% of cells). The mixed suspension with nuclei from both patients is analysed using sci-HIPSD-seq (left heatmap) and the 1% clone can be robustly identified by correlating CNVs from every cell to the two bulk profiles (bottom right). Additionally, we also performed HIPSD-seq for patient 41 (grey dots) that aligns to the major clone of the sci-HIPSD-seq profile.

## Discussion

We have established HIPSD&R-seq, a new single-cell DNA sequencing assay, which (i) allows copy-number profiling in thousands of cells, (ii) is compatible with multiome read-out, and (iii) offers sparse but uniform coverage (**Suppl. Fig. 1**). We provide computational workflows that are tailored to analyse sparse data, borrowing concepts from meta celling that were established for scRNA-seq. HIPSD&R-seq can be applied to a range of sample types, is easily implemented and thus widely applicable. We showed that HIPSD&R-seq DNA provides a significant improvement in CNV resolution compared to inference-based methods applied to the RNA component of the same cells, and yields results that are markedly concordant with CNVs detected in matched bulk WGS samples. When combining HIPSD-seq with previously proposed combinatorial indexing to increase the throughput further, we improve the coverage and reach an ultra-high throughput, recovering a subclone making up 1% of the cell population and identifying major clones with distinct CNV profiles.

For the multiome assay, one limitation of the method is the high duplication rate, which is due to the lower activity of the transposase of the 10X multiome kit, as compared to the highly concentrated Tn5 that ensures high integration rates across the genome used in the HIPSD-seq assay. However, HIPSD&R-seq provides CNV profiles at high resolution and matched transcriptomes for the same cells, opening up a number of applications in cancer research and beyond. As compared to methods such as DLP+, which provide excellent CNV read-outs but require specialised equipment available to very few laboratories in the world, HIPSD-seq can be implemented anywhere. The high yield will lead to a better understanding of genetic heterogeneity and chromosome instability in space and time, by enabling high-throughput analyses of longitudinal and multi-region sampling.

## Conclusion

Dissecting matched single-cell genomes and transcriptomes will lead to a better understanding of the transcriptional consequences of genetic variation and allow to disentangle the links between genome and transcriptome. Ultimately, capturing CNV-based cell lineage trees with cell type and state annotations will help to decipher the role of cell heterogeneity in disease and evolution.

## Methods

### Tn5 loading

Lyophilized adapter oligos were resuspended at 100 μM in annealing buffer (50 mM NaCl, 40 mM Tris pH 8). Adapters (Oligo list, see Supplementary Table 2) were pre-annealed on a thermocycler for 2 min at 85°C followed by cooling down to 20 °C with one °C per minute. For HIPSD-seq, Read1 was annealed with blocked-phos-ME and Read2 with blocked-phos-ME. For sci-HIPSD-seq, Read1 was annealed with unblocked-phos-ME and barcoded-Read2 with unblocked-phos-ME. Equal volume of 100% glycerol was then added to the adapters and they were stored at −20°C. For Tn5 assembly, transposase and annealed primers are mixed with equal volumes and incubated for 30 min at room temperature. The Tn5 was then diluted in dilution buffer (50 mM Tris pH 7.5, 100 mM NaCl, 0.1 mM EDTA, 1 mM DTT, 0.1% NP-40, 50% Glycerol) to 83 µg/ml for HIPSD-seq.

### Cell culture and nuclei extraction from fibroblasts

#### Sample preparation for fibroblasts with high genome instability

Fibroblasts from patients with Li Fraumeni Syndrome (germline variant in *TP53*) were grown in Minimum Essential Medium Eagle (Sigma; M5650) supplemented with 10% foetal calf serum, 1% glutamine and 1% penicillin/streptomycin and cultured at 37°C with 5% CO2. For mATAC, cells from LFS041 p.27 and LFS041 p.63 were harvested after trypsinizing the cells with 0.25% trypsin and resuspending them in PBS. For scMultiome, cells from LFS041 p.22 and LFS041 p.63 were harvested in the same way. For sciDNA-seq, cells from LFS041 p.63 and LFS087 p.195 were harvested in the same way, and resuspended in PBS with 1% BSA.

#### Nuclei extraction and nucleosome depletion for fibroblasts

For HIPSD-seq, 1 × 10^6^ cells were fixed in 1 mL of 1.5% methanol-free formaldehyde (FA; Thermo Scientific™, #28906) in PBS, for 10 min at room temperature with gentle shaking. Fixation was neutralized by adding 200 mM Glycine, followed by incubation on ice for 5 min. Cells were centrifuged (550xg, 5 min, 4°C) and washed with ice-cold PBS. For nuclei isolation, cells were resuspended in 1 mL of ice-cold NIB buffer (10 mM TrisHCl pH7.4, 10 mM NaCl, 3 mM MgCl2, 0.1% igepal, 1x protease inhibitor cocktail (#5871S, Cell Signaling Technology)) and incubated on ice for 20 min with gentle mixing. Nuclei were centrifuged (500xg, 5 min, 4°C) and washed once with 1X NEBuffer 2.1 (NEB, #B7202). For nucleosome depletion, nuclei were resuspended in 1X NEBuffer 2.1 supplemented with 0.3% SDS (Serva, #20767) and incubated at 42°C, for 15 min with shaking. SDS was quenched by adding 2% Triton-X100 (Sigma-Aldrich, #93443), followed by incubation at 42°C, for 15 min with shaking. Nuclei were centrifuged (500xg, 5 min, 4°C) and resuspended in 1X nuclei buffer (10X Genomics). Nuclei were counted on the Luna-FL^TM^ cell counter (Logos biosystems) and diluted to 2000-5000 nuclei /μL.

For HIPSD&R-seq, fixation with 1.5% FA, nuclei isolation with NIB buffer and nucleosome depletion with 0.3% SDS were performed exactly as in case of HIPSD-seq, with the exception that 1U/μL of RNAse inhibitor (Takara Bio, #2313A) was added to all buffers during and after nuclei isolation (i.e. to NIB buffer and 1X NEBuffer 2.1). Following quenching of nucleosome depletion with 2% Triton-X100, nuclei samples were centrifuged (500xg, 5 mins, 4°C) and resuspended in 1X Nuclei Buffer (10X Genomics) containing 1U/μL of RNAse inhibitor. Nuclei were counted using the Luna-FL^TM^ cell counter and diluted to 2000-5000 nuclei/μL.

For sci-HIPSD-seq, 1.66 × 10^6^ cells (LFS041 p.63) and 2.7 × 10^5^ cells (LFS087 p.195) were used. Fixation with 1.5% FA, nuclei isolation with NIB buffer and nucleosome depletion with 0.3% SDS were performed exactly as in case of HIPSD-seq, with the exception that all the centrifugation steps were performed at 600xg for 8 min at 4°C. Following quenching of nucleosome depletion with 2% Triton-X100, nuclei samples were centrifuged (600xg, 8 min, 4°C) and resuspended in in ice-cold basic buffer (1 mM DTT, 2%BSA in 1xPBS). Nuclei were counted on the Luna-FL^TM^ cell counter. 600,000 nuclei of LFS041 p.63 were used for transposition. 6,000 nuclei of LFS087 p.195 were spiked-in during transposition.

### HIPSD-seq from fibroblasts

1 × 10^6^ cells (LFS041 p.27, LFS041 p.63) were used for nuclei extraction and nucleosome depletion as described above. The nucleosome-depleted nuclei were processed using the 10X ATAC protocol, according to the Chromium Single Cell ATAC Reagent Kits User Guide v1.1 Chemistry (CG000209, 10X Genomics), but the transposase provided by 10X Genomics was replaced by highly active in-house Tn5 (83 µg/ml). 10,000 nuclei were used for transposition and loaded on the Chromium Next GEM Chip H (PN-1000161). scDNA libraries were generated using standard reagents from the Chromium Next GEM Single Cell ATAC Library & Gel Bead Kit v1.1, (PN-1000176). Libraries were uniquely indexed with Single Index Kit N Set A (PN-1000212). Quality control and molarity calculations of the final libraries were performed with the Qubit 3.0 Fluorometer (#Q33216, Invitrogen) and the 4200 Tapestation system (Agilent Technologies). Libraries were sequenced using NovaSeq 6000 paired-end 100 SP with read 1 sequenced for 51 cycles read 2 for 51 cycles, 8 cycles for i7, and 16 cycles for index 2. Libraries were sequenced with 1% PhiX.

### HIPSD&R-seq from fibroblasts

1 × 10^6^ cells (LFS041 p.22, LFS041 p.63) were used for nuclei extraction and nucleosome depletion as described earlier. The nucleosome depleted nuclei were processed using the 10X Multiome protocol, according to the Chromium Next GEM Single Cell Multiome ATAC + Gene Expression Reagent Kits User Guide (CG000338, 10X Genomics). 10,000 nuclei were used for transposition and loaded on the Chromium Next GEM Chip J (PN-1000230). scDNA and scRNA libraries were generated using the standard reagents provided in the Chromium Next GEM Single Cell Multiome ATAC + Gene Expression Reagent Bundle, (PN-1000285). scDNA libraries were indexed using Single Index Kit N Set A, (PN-1000212), while the scRNA libraries were indexed with Dual Index Kit TT Set A, (PN-1000215). Quality control and molarity calculations of the final libraries were performed with the Qubit 3.0 Fluorometer (#Q33216, Invitrogen) and the 4200 Tapestation system (Agilent Technologies). scDNA libraries were sequenced using NovaSeq 6000 (100 cycles) S1. 51 cycles were used for read 1, 51 cycles for read 2, 8 cycles for i7, and 24 cycles for i5. scRNA libraries were sequenced using NovaSeq 6000 paired-end 100 SP with read 1 for 28 cycles, read 2 for 90 cycles, i7 for 10 cycles and i5 for 10 cycles. Libraries were sequenced with 1% PhiX.

### sci-HIPSD-seq protocol with 1% spike-in (skin-derived fibroblasts)

1.66 × 10^6^ cells (LFS041 p.63) and 2.7 × 105 cells (LFS087 p.195) were used for nuclei extraction and nucleosome depletion as described earlier.

#### Tn5 assembly

1.1 μl of Tn5-Read1 was mixed with 1.1 μl of each Tn5-Barcoded Read2 into 30 individual wells of a 96-well plate. For 600,000 nuclei, we used 30 Read2 barcodes (Oligo list, see Supplementary Table 2). One μl of barcoded annealed oligos were mixed with 1 μl of highly active in-house Tn5 into 30 wells of a new 96-well plate and Tn5 assembly was performed for 30 min at room temperature.

#### Transposition

For transposition, 2X TD was prepared by mixing equal volumes of 100% DMF and 4x Tn5 buffer (40 mM Tris-HCl pH 7.5, 40 mM MgCl2). 600,000 nuclei (LFS041 p.63) and 6,000 nuclei (LFS087 p.195) were used for transposition in 1X TD buffer [83]. Roughly, 20,200 nuclei (18 μL of transposition mix) were added to each of the 30 wells of the 96-well plate containing 2 μL of assembled Tn5 (final dilution of Tn5 in the reaction is 1:10). The plate was incubated at 37°C for 60 min with shaking at 500 rpm and then transferred to ice. 1x PBS with 1% BSA (40 μL) was added to each of the wells and the samples were pooled together into Eppendorf tubes. Wells were washed with additional 40 μL of 1x PBS with 1% BSA and transferred to the same tubes. Nuclei were collected by centrifugation (500xg, 8 min, 4°C) and resuspended in 500 μL of 1x PBS with 2% BSA. This nuclei suspension was filtered through a pre-wet 30 μm filter (Militenyi Biotec, #130-041-407) to avoid clogging of the 10X chip. Filter was rinsed once with 500 μL of 1x PBS with 0.5% BSA and collected into the sample tube. Nuclei were pelleted by centrifugation (500xg, 5 min, 4°C), resuspended in 20 μL of 1x Nuclei Buffer (10X Genomics) and counted on Luna-FL^TM^ cell counter.

#### sci-HIPSD Library preparation and sci-HIPSD-seq

This nuclei suspension was then processed using the 10X ATAC protocol, according to the Chromium Single Cell ATAC Reagent Kits User Guide v1.1 Chemistry (CG000209, 10X Genomics). 180,000 nuclei were mixed with the barcoding reaction mix and loaded on the Chromium Next GEM Chip H (PN-1000161) for GEM formation. sci-HIPSD libraries were generated using standard reagents from the Chromium Next GEM Single Cell ATAC Library & Gel Bead Kit v1.1, (PN-1000176). Following the clean-up with 1.2x AMPure XP Beads, the sample was pre-amplified using Partial P5 and Partial P7 primers (Oligo list, see Supplementary Table 2). 2.5 μL of the pre-amplified sample was used for qPCR to determine the additional number of cycles required for library amplification. The sciDNA library was then amplified using the optimal number of cycles determined from qPCR and a double size selection was performed with a 0.6x and 1.2X ratio of AMPure XP beads (#A63882, Beckman Coulter). Quality control and molarity calculations of the final libraries were performed with the Qubit^ä^ 3.0 Fluorometer (#Q33216, Invitrogen) and the 4200 Tapestation system (Agilent Technologies).

sci-HIPSD libraries were sequenced using NovaSeq 6000 paired-end 100 S1 v1.5. 300 pM was loaded and 101 cycles were used for read 1, 101 cycles for read 2, 11 cycles for i7, and 16 cycles for i5. Libraries were sequenced with 1% PhiX.

### Bulk whole-genome sequencing for skin-derived fibroblasts

Cell pellets from fibroblasts (LFS041, passage 63 and LFS087, passage 195) were prepared after trypsinization in 0.25% trypsin, cell resuspension and centrifugation. The cell pellets were kept at −80°C until performing the DNA extraction using DNAeasy Blood & Tissue Kit (QIAGEN; Cat. No. ID: 69504). Qubit Broad Range double-stranded DNA assay (Life Technologies) was used to quantify the DNA, while a Bioanalyzer (Agilent) was used to measure the quality of the DNA. Whole genome sequencing was done using the Illumina NovaSeq 6K paired-end 150 SP platform.

### HIPSD&R-seq in patient-derived xenograft (PDX) cells

Cryopreserved PDX cells (passage 1, primary tumour from a patient with Sonic Hedgehog medulloblastoma with germline variant in *TP53*) were thawed in a 37°C water bath and suspended in high purity grade PBS supplemented with 10% foetal calf serum. Following suspension, the cells were centrifuged at 1000 rpm for 5 min at 4°C and washed two times to remove any remaining DMSO. The cells were filtered with 40 µm cell strainer and brought to FACS sort in 2 ml of supplemented PBS. Directly before the FACS sort, the cells were stained with TO-PRO-3 in order to exclude any dead cells. Contaminating mouse cells were excluded based on the side forward scatter.

### Nucleosome depletion and HIPSD&R-seq library preparation (PDX cells)

420,000 FACS isolated PDX cells were used. For HIPSD&R sample preparation, fixation with 1.5% FA, nuclei isolation and nucleosome depletion were performed exactly as in case of HIPSD&R for fibroblasts, with the exception that all centrifugation steps were performed at 800xg for 5 min at 4°C. Following quenching of nucleosome depletion with 2% Triton-X100, nuclei were centrifuged (800xg, 5 min, 4°C) and resuspended in 1x Nuclei Buffer (10X Genomics) with 1U/μL RNase inhibitor to a final concentration of 2000-5000 nuclei/μL. 10,000 nuclei were used for transposition and HIPSD&R library preparation following the exact 10X Multiome protocol mentioned in case of HIPSD&R from fibroblasts.

scDNA libraries were sequenced using NovaSeq 6000 (138 cycles) SP, with 51 cycles for Read 1, 51 cycles for Read 2, 8 cycles for index 1, and 24 cycles for index 2. scRNA libraries were sequenced using NovaSeq 6000 (138 cycles) SP, with 28 cycles for Read 1, 90 cycles for Read 2, 10 cycles for i7 and 10 cycles for i5. Libraries were sequenced with 1% PhiX.

### 10X CNV single-cell DNA-sequencing library preparation (PDX cells)

The single-cell suspensions from PDX cells were processed using the Chromium Single-Cell CNV Kit (10× Genomics) [81] according to the manufacturer’s protocol. In brief, using cell bead polymer, single cells were partitioned in a hydrogel matrix on Chromium Chip C. Once the cell beads were encapsulated and incubated, they were subjected to enzymatic and chemical treatment. This lysed the encapsulated cells and denatured the gDNA in the cell bead, to make it accessible for further amplification and barcoding. A second encapsulation was performed to achieve single cell resolution by co-encapsulating a single cell bead and a single barcoded gel bead to generate GEMs. Immediately after GEM generation the gel bead and cell bead were dissolved. Oligonucleotides containing standard Illumina adaptors and 10x barcoded fragments were then amplified with 14 PCR cycles during two-step isothermal incubation. After incubation the GEMs were broken and pooled 10x barcoded fragments were recovered. Single-cell libraries were sequenced on the Illumina NextSeq platform (150 bp, paired-end). For more details see Parra et al. [82].

### Computational analyses

#### HIPSD-seq raw data preprocessing

The initial preprocessing of the samples is performed using the 10x Cellranger ATAC pipeline [84]. To avoid defining cells based on peak calling results, we define cells based on number of fragments per cell and uniformity of coverage. The count cut-off is defined based on the elbow plot.

When the cell barcodes are retrieved, we extract cell-specific BAM files from Cellranger produced BAM file using subset-bam [85] tool from 10x Genomics. Individual BAM files are preprocessed by removing low quality reads and duplicates (using SAMtools [86]) and then counts are extracted using hmmcopy_utils at 100kb resolution [87].

#### Metacelling

To perform high confidence copy number calling, we aggregate cells into metacells (**Supp. Fig. 2**). The aggregation is performed in the following steps:

1. Cell counts are aggregated into 1MB windows to increase signal-to-noise ratio and reduce sparsity.
2. Principal components are computed to reduce dimensionality, while preserving 95% of variance.
3. Cells are preclustered using leiden clustering (based on the principal components) such that the maximal number of reads per 1MB does not exceed 1200 (can be selected by user). This step creates initial metacells that should be fine-grain enough to avoid clamping together genetically distinct cells.
4. Premetacells are separated into three categories: those with insufficient coverage (selected by user - 200 reads per 1MB in our case that matches 0.02x coverage), those with sufficient coverage, those with high coverage (over 1200 reads per 1MB).
5. A greedy algorithm ranks every cell pair by genetic distance and makes a full pass merging premetacells within fixed distance (selected by a user) if both cells have insufficient coverage, or one cell has insufficient coverage and one sufficient. Premetacells with high coverage are not merged with any other cells as we want to avoid creating cells with a coverage that is way higher than others.
6. Steps 4 and 5 are repeated until there are no more possible merges.

Selecting appropriate merging distance for step 5 is essential. The value is chosed such that the trade-off between within metacell variation (high with large distances) and unmerged cells (high with small distances) is optimal. In our case we computed 10 quantiles of all distances between premetacells and picked as the final merging distance the one that gives the best within metacell variation vs. number of unmerged cells trade-off. More precisely the value for HIPSD-seq was 34.97, for sciHIPSD-seq 34.11 and for HIPSD&R-seq 46.48.

**Supp. Fig. 2** shows the metacelling steps and results using as an example HIPSD-seq sample from LFS041 p.63. The code is available at https://github.com/Laolga/HIPSD_seq [88].

#### Copy number calling for scDNA-seq data

For DNA-seq data (HIPSD-seq, sciHIPSD-seq or WGS) we called copy number profiles using HMMCopy tool [89]. For single cell approaches we applied metacelling first to increase the coverage per cell (see **Suppl. Fig. 1 G** for pre-metacelling coverage and **Suppl. Fig. 1 A** for post-metacelling coverage, as well as **Suppl. Fig. 1 E** too see cell per metacell assigment). HMMCopy was set up to produce long fragments. We noticed the tendency of HMMCopy to overcall homozygous deletions for all datasets regardless of coverage and therefore we do not distinguish homozygous and heterozygous deletions in our results.

#### scRNA data analysis

scRNA data was processed using the Cellranger pipeline [65]. We used Numbat [47] with default parameters to estimate copy numbers from scRNA-seq data.

## Declarations

### Ethics approval and consent to participate

Not applicable.

### Consent for publication

Not applicable.

### Availability of data and materials

All raw sequencing data for this study will be available at the European Genome-Phenome Archive (EGA) under accession number <> (deposition is in progress). Before the deposition, the data can also be requested from corresponding authors upon request.

10x Genomics CNV and RNA data is obtained from Parra et al. [82] and available from EGA under accession number EGAS00001005410.

The code to reproduce presented results is available at GitHub repository https://github.com/Laolga/HIPSD_seq [88].

## Authors’ contributions

OL did the vast majority of the computational analyses and generated figures. J-P M conceptualised and supervised the experimental work and performed part of it. MS performed the xenograft experiments. GP performed the experiments with fibroblasts. PS supported the nuclei extractions and did the library preparations. UP and LH supported the experimental work. AL supported the computational analyses. UY and MM supported the combinatorial indexing experiments. JZ and KMN contributed to conceptualization and interpretation. OS and AE conceived the study and jointly supervised the work. AE acquired the financial support for the project. All authors discussed the results and contributed to the final manuscript.

## Acknowledgements

We thank Frauke Devens, Michaela Hergt and Michèle Vousten for excellent technical support. We thank the Genomics Core Facility of the German Cancer Research Center. We thank Kim Remans and her team from the EMBL Protein Expression and Purification Core Facility for providing the Tn5.

## Additional files

File name: supplement.pdf. File format: PDF file.

Title of data: Supplementary information.

Description of data: Contains Supplementary Table 1 and 2 and Supplementary Figures 1 and 2.

