## Supplementary material for "HIPSD&R-seq enables scalable genomic copy number and transcriptome profiling"

**Supplementary Table 1:** Samples characteristics. Table provides the list of samples used in the paper and the key QC metrics including tissue description; passage (applicable only to fibroblast samples); assay used; number of cells obtained based on elbow-plot cut-off for DNA data or based on cellranger definition for RNA data; as well as median counts per cell, genes per cell, duplication and sequencing saturation.

| Tissue/ sample type | Short sample name | Passage | Assay Types | # cells (DNA/RNA/ shared) | DNA | | RNA | |
|---|---|---|---|---|---|---|---|---|
| | | | | | Median counts per cell | Median percent duplicates | Median genes per cell | Sequencing Saturation |
| Fibroblasts from LFS patient 41, passage 63 | LFS041_63 | 63 | HIPSD-seq | 5,084 | 99,930 | 18% | - | - |
| Fibroblasts from LFS patient 41, passage 63, with a 1% spike-in from patient 87 | LFS041_63_ spikein | 63 | sciHIPSD-seq | 17,929 | 18,798 | 25% | - | - |
| Patient-derived xenograft of medulloblast | PDX | - | HIPSD&R-seq | 2,658/2687/ 2597 | 137,927 | 91% | 1,870 | 96.2% |

| | | | | | | | | |
|---|---|---|---|---|---|---|---|---|
| oma | | | | | | | | |
| Patient-derived xenograft of medulloblastoma | PDX | - | Chromium Single Cell CNV | 244 | 1,832,718 | 34% | - | - |
| Patient-derived xenograft of medulloblastoma | PDX | - | Chromium Single Cell Gene Expression | 5012 | - | - | 1,071 | 96.8% |

**Supplementary Table 2:** List of oligos used in the HIPSD-seq and sci-HIPSD-seq assays

| Oligo name | Oligo sequence | Assay |
|---|---|---|
| Read1 | 5′-TCGTCGGCAGCGTCAGATGTGTATAAGAGACAG | HIPSD-seq, sci-HIPSD-seq |
| phos-Read2 | 5′-[phos]GTCTCGTGGGCTCGGAGATGTGTATAAGAGACAG | HIPSD-seq |
| Blocked phos-ME | 5'-[Phos]C*T*G*T*C*T*C*T*T*A*T*A*C*A*[23ddC] | HIPSD-seq |
| Unblocked phos-ME | 5'- [Phos]CTGTCTCTTATACACATCT | sci-HIPSD-seq |
| sciDNA_ barcoded Read2 | 5′-CAAGCAGAAGACGGCATACGAGAT[round1_index]GTCTCGTGGGCTCGGAGATGTGTATAAGAGACAG | sci-HIPSD-seq |
| Barcodes used: | | |
| sciDNA_001 | 5' -CAAGCAGAAGACGGCATACGAGATACCTCCAACCCGTCTCGTGGGCTCGGAGATGTGTATAAGAGACAG | sci-HIPSD-seq |
| sciDNA_002 | CAAGCAGAAGACGGCATACGAGATCGTTTCAGCCAGTCTCGTGGGCTCGGAGATGTGTATAAGAGACAG | sci-HIPSD-seq |
| sciDNA_003 | CAAGCAGAAGACGGCATACGAGATTAAGGTTAGATGTCTCGTGGGCTCGGAGATGTGTATAAGAGACAG | sci-HIPSD-seq |
| sciDNA_004 | CAAGCAGAAGACGGCATACGAGATATATACCAGACGTCTCGTGGGCTCGGAGATGTGTATAAGAGACAG | sci-HIPSD-seq |
| sciDNA_005 | CAAGCAGAAGACGGCATACGAGATGGAGAGACTGTGTCTCGTGGGCTCGGAGATGTGTATAAGAGACAG | sci-HIPSD-seq |
| sciDNA_006 | CAAGCAGAAGACGGCATACGAGATCTCATCTTAGCGTCTCGTGGGCTCGGAGATGTGTATAAGAGACAG | sci-HIPSD-seq |
| sciDNA_007 | CAAGCAGAAGACGGCATACGAGATAATTTGGCCGTGTCTCGTGGGCTCGGAGATGTGTATAAGAGACAG | sci-HIPSD-seq |

| | | |
|---|---|---|
| sciDNA_008 | CAAGCAGAAGACGGCATACGAGATTTAAGGAAGTAGTCTCGTGGGCTCGGAG ATGTGTATAAGAGACAG | sci-HIPSD-seq |
| sciDNA_009 | CAAGCAGAAGACGGCATACGAGATACCGATCCAGAGTCTCGTGGGCTCGGAG ATGTGTATAAGAGACAG | sci-HIPSD-seq |
| sciDNA_010 | CAAGCAGAAGACGGCATACGAGATCAACGGGACTGGTCTCGTGGGCTCGGAG ATGTGTATAAGAGACAG | sci-HIPSD-seq |
| sciDNA_011 | CAAGCAGAAGACGGCATACGAGATTCCAGTACAACGTCTCGTGGGCTCGGAG ATGTGTATAAGAGACAG | sci-HIPSD-seq |
| sciDNA_012 | CAAGCAGAAGACGGCATACGAGATGAGGACGATGCGTCTCGTGGGCTCGGAG ATGTGTATAAGAGACAG | sci-HIPSD-seq |
| sciDNA_013 | CAAGCAGAAGACGGCATACGAGATAAGGGTCGCTTGTCTCGTGGGCTCGGAG ATGTGTATAAGAGACAG | sci-HIPSD-seq |
| sciDNA_014 | CAAGCAGAAGACGGCATACGAGATCCCATGTAGCAGTCTCGTGGGCTCGGAG ATGTGTATAAGAGACAG | sci-HIPSD-seq |
| sciDNA_015 | CAAGCAGAAGACGGCATACGAGATGTGAAGTGGACGTCTCGTGGGCTCGGAG ATGTGTATAAGAGACAG | sci-HIPSD-seq |
| sciDNA_016 | CAAGCAGAAGACGGCATACGAGATTAATAATTACAGTCTCGTGGGCTCGGAGA TGTGTATAAGAGACAG | sci-HIPSD-seq |
| sciDNA_017 | CAAGCAGAAGACGGCATACGAGATGCTGTAACGTGGTCTCGTGGGCTCGGAG ATGTGTATAAGAGACAG | sci-HIPSD-seq |
| sciDNA_018 | CAAGCAGAAGACGGCATACGAGATAAAAGCTGCGTGTCTCGTGGGCTCGGAG ATGTGTATAAGAGACAG | sci-HIPSD-seq |
| sciDNA_019 | CAAGCAGAAGACGGCATACGAGATCGCGCAAACAGGTCTCGTGGGCTCGGAG ATGTGTATAAGAGACAG | sci-HIPSD-seq |
| sciDNA_020 | CAAGCAGAAGACGGCATACGAGATTAATTTTCCCCGTCTCGTGGGCTCGGAGA TGTGTATAAGAGACAG | sci-HIPSD-seq |
| sciDNA_021 | CAAGCAGAAGACGGCATACGAGATGGATACTGGTTGTCTCGTGGGCTCGGAG ATGTGTATAAGAGACAG | sci-HIPSD-seq |
| sciDNA_022 | CAAGCAGAAGACGGCATACGAGATAGGGGGAAGCGGTCTCGTGGGCTCGGA GATGTGTATAAGAGACAG | sci-HIPSD-seq |
| sciDNA_023 | CAAGCAGAAGACGGCATACGAGATCAACGCCAAAAGTCTCGTGGGCTCGGAG ATGTGTATAAGAGACAG | sci-HIPSD-seq |
| sciDNA_024 | CAAGCAGAAGACGGCATACGAGATACGAAGCCTGTGTCTCGTGGGCTCGGAG ATGTGTATAAGAGACAG | sci-HIPSD-seq |
| sciDNA_025 | CAAGCAGAAGACGGCATACGAGATTACGTTGCCAGGTCTCGTGGGCTCGGAG ATGTGTATAAGAGACAG | sci-HIPSD-seq |
| sciDNA_026 | CAAGCAGAAGACGGCATACGAGATAATTAACTTATGTCTCGTGGGCTCGGAGA TGTGTATAAGAGACAG | sci-HIPSD-seq |
| sciDNA_027 | CAAGCAGAAGACGGCATACGAGATCTCGGTATGGGGTCTCGTGGGCTCGGAG ATGTGTATAAGAGACAG | sci-HIPSD-seq |
| sciDNA_028 | CAAGCAGAAGACGGCATACGAGATTCGTGTGAATTGTCTCGTGGGCTCGGAGA TGTGTATAAGAGACAG | sci-HIPSD-seq |
| sciDNA_029 | CAAGCAGAAGACGGCATACGAGATATAATTAGTTCGTCTCGTGGGCTCGGAGA TGTGTATAAGAGACAG | sci-HIPSD-seq |
| sciDNA_030 | CAAGCAGAAGACGGCATACGAGATGGCTAATTCCTGTCTCGTGGGCTCGGAG ATGTGTATAAGAGACAG | sci-HIPSD-seq |
| Partial P5 | 5‘ - AATGATACGGCGACCACCGAG*A | sci-HIPSD-seq |
| Partial P7 | 5‘ - CAAGCAGAAGACGGCATACGA*G | sci-HIPSD-seq |

Abbreviations: phos, 5’ phosphorylation; *, PTO modification

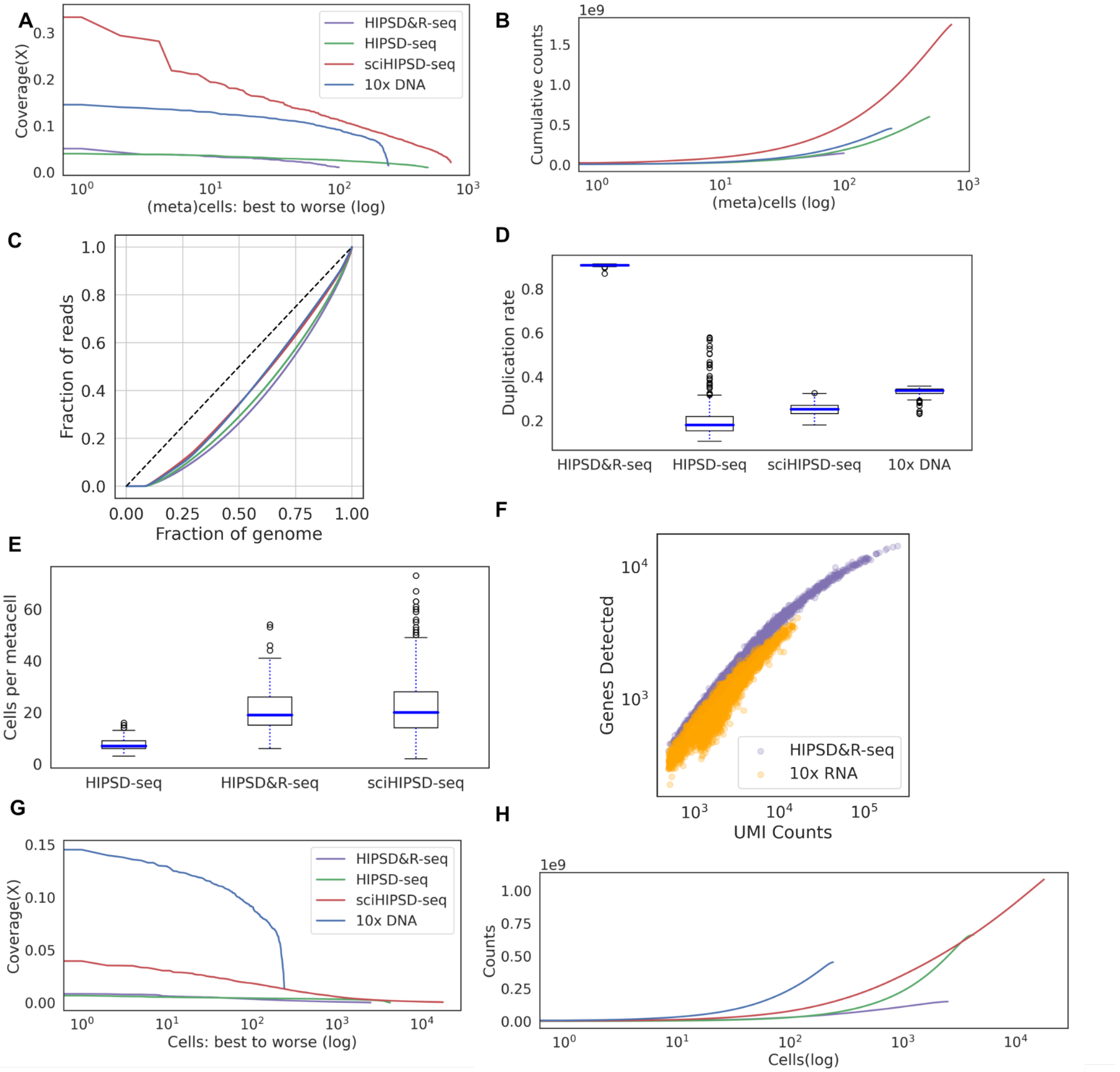

**Supp. Figure 1 | Assay properties and qualitry metrics on metacells and cells (HIPSD&R-seq - PDX sample, HIPSD-seq - LFS041_63 sample, sci-HIPSD-seq - LFS041_63_spikein sample) in comparison to Chromium Single Cell CNV - PDX sample.** **(A)** Read coverage of processed and aggregated data for HIPSD&R-seq/HIPSD-seq/sciHIPSD-seq (metacells), and for data from the 10x CNV kit (individual cells). (**B)** Cumulative read counts per metacell for HIPSD&R-seq/HIPSD-seq/sciHIPSD-seq and per cell for the 10x CNV kit. (**C)** Lorenz curves for the same assays as in **A**. **(D)** Duplication rate for the same assays as in **A**. (**E)** Statistics for metacell definitions for the HIPSD&R-seq assays as in **A**. (**F**) Sequence saturation for the HIPSD&R-seq RNA component versus conventional 10x 3' scRNA-seq on the original cells of the PDX sample.(**G)** Coverage without metacelling. (**H)** Cumulative counts distribution without metacelling.

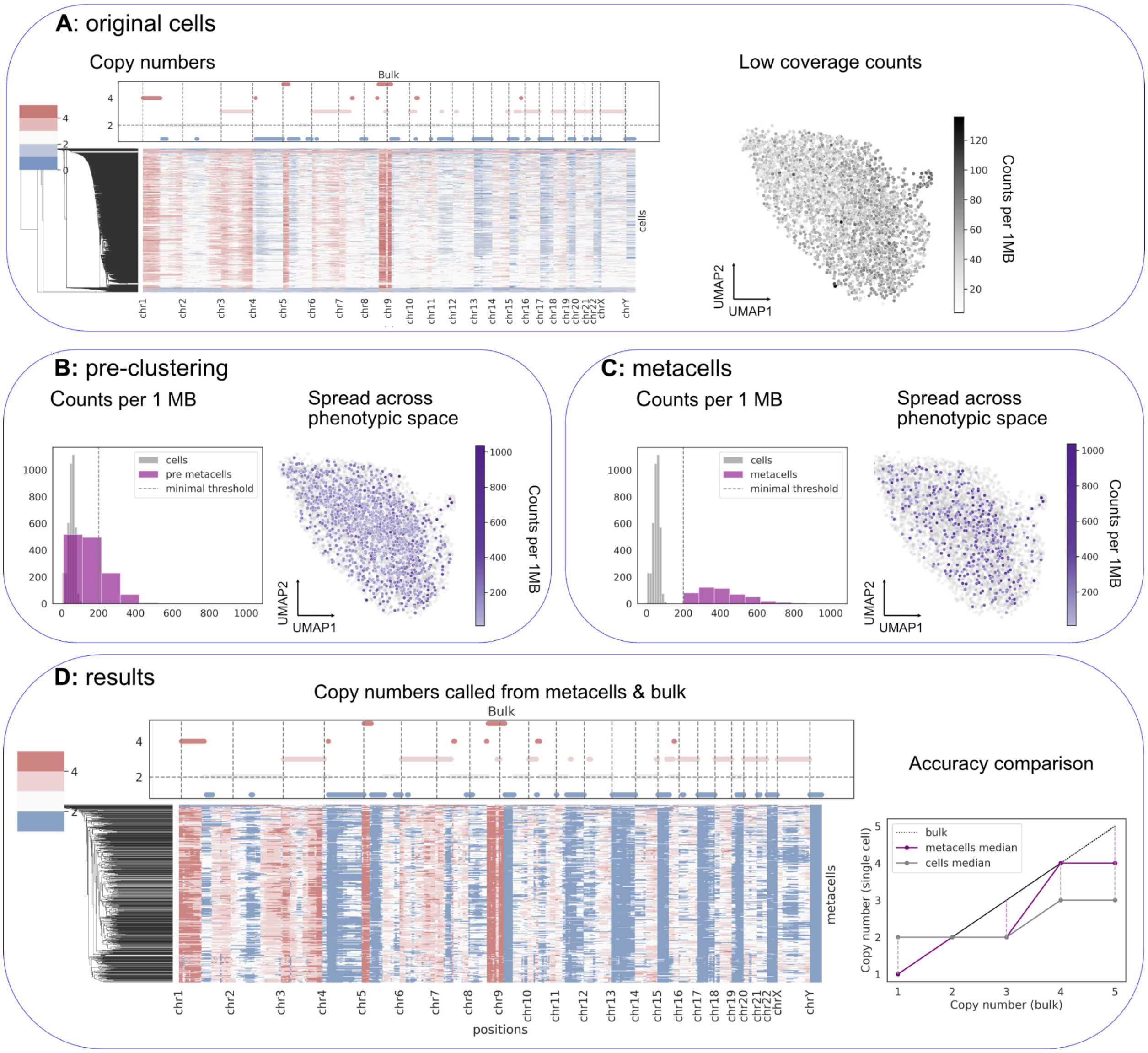

**Supp. Figure 2 | Metacelling workflow. (A)** Pre metacelling results. Left: copy number estimates on 100kb resolution for bulk data (top) and single-cell (bottom); right: counts UMAP for 1MB window. (**B)** Cells are preclustered to identify candidate metacells, where some of the clusters might not have sufficient coverage. (**C)** Pre-metacells are greedily merged within a fixed distance until no metacell has a coverage lower than a selected threshold. (**D)** Copy number called on metacells (bottom) and corresponding bulk with fewer mismatches (top); comparison between bulk, initial cells and metacells shows that metacells produce closer match to bulk.
